# Improving range shift predictions: enhancing the power of traits

**DOI:** 10.1101/2021.02.15.431292

**Authors:** Anthony F. Cannistra, Lauren B. Buckley

**Author notes:** Corresponding author: Anthony F. Cannistra.

## Abstract

Accurately predicting species’ range shifts in response to environmental change is a central ecological objective and applied imperative. In synthetic analyses, traits emerge as significant but weak predictors of species’ range shifts across recent climate change. These studies assume linearity in the relationship between a trait and its function, while detailed empirical work often reveals unimodal relationships, thresholds, and other nonlinearities in many trait-function relationships. We hypothesize that the use of linear modeling approaches fails to capture these nonlinearities and therefore may be under-powering traits to predict range shifts. We evaluate the predictive performance of four different machine learning approaches that can capture nonlinear relationships (ridge-regularized linear regression, ridge-regularized kernel regression, support vector regression, and random forests). We validate our models using four multi-decadal range shift datasets in montane plants, montane small mammals, and marine fish. We show that nonlinear approaches perform substantially better than least-squares linear modeling in reproducing historical range shifts. In addition, using novel model observation and interrogation techniques, the trait classes (e.g. dispersal-or diet-related traits) that we identify as primary drivers of model predictions are consistent with expectations. However, disagreements among models in the directionality of trait predictors suggests limits to trait-based statistical predictive frameworks.

## 1. INTRODUCTION

Species have been responding to recent climate change by tracking their environment in space or time, adapting or acclimating, or facing declines (Parmesan, 2006), but we are largely unable to predict how particular species will respond (Maguire et al., 2015). Extensive documentation of shifts in distribution and seasonal timing (phenology) reveal that responses vary among species markedly in direction and extent (Rapacciuolo et al., 2014). Detailed empirical studies often succeed in identifying functional traits that govern climate change responses (e.g., Adrian et al., 2006) and consequently functional ecology has been rapidly gaining prominence in climate change ecology (Buckley & Kingsolver, 2012). However, attempts to use traits to predict the relative magnitude of responses among species generally identify traits that are significant, but weak (accounting for ~16% of the among species variation in range shifts, Buckley & Kingsolver, 2012), predictors of climate change responses (Estrada et al., 2016; MacLean & Beissinger, 2017). How can we close the discrepancy between traits predicting responses well in detailed studies but poorly in broad studies? What statistical techniques will allow us to generalize the importance of traits in mediating climate change responses?

Addressing such questions is imperative for anticipating and adapting to the biological impacts of climate change. Indeed, traits are already being used to predict species’ sensitivity to climate change in vulnerability frameworks (Foden et al., 2013). However, the frameworks remain largely untested and perform poorly in tests (Wheatley et al., 2017). Most attempts to use species’ traits to predict the magnitude of their climate change responses rely on linear regression (Buckley & Kingsolver, 2012; MacLean & Beissinger, 2017), yet detailed empirical studies often reveal non-linear relationships between traits and their function (Stenseth & Mysterud, 2002). Unimodal relationships and thresholds are common. For example, extreme diet specialization may drive a species to track the range shift of a food item (Diamond et al., 2011), but reducing diet specialization only slightly may alleviate the need for a species to track its food. Diet generalization could facilitate species moving to capitalize on newly climatically suitable habitat, yielding a unimodal relationship between diet specialization and the magnitude of range shifts. Likewise, low dispersal ability may prevent a species from tracking its environmental niche (Schloss et al., 2012), but the threshold of dispersal ability that allows species to track their niche may be relatively low. Trade-offs among traits and differences in the developmental dependencies of traits may also produce nonlinearities (Fitt et al., 2018). Can statistical techniques that allow for non-linear relationships between traits and species’ responses improve our predictive ability?

Non-linear modelling techniques have been underutilized in predicting climate change responses (Olden et al. 2008). Standard approaches to capture variable interactions and nonlinearities in linear regressions (such as the explicit inclusion of interacting variables or polynomial expansion) rely on prior knowledge or model selection techniques to determine which variables to select. Other model types, such as machine learning approaches that optimize model parameters, are better suited to capture functional relationships among variables. These models, while offering statistical-robustness and efficiency, can be opaque and rarely afford clear coefficients to inspect when assessing the model’s learned correlations. However, machine learning developments offer new approaches for inspecting model performance and predictions.

Here we assess whether machine learning-based models can better use species’ traits to predict the magnitude and direction of range shifts observed in response to recent climate change. First, we consider whether several models which are able to capture nonlinear relationships can outperform linear models in their predictive ability. Second, we use recently developed model interpretability techniques to ask whether model predictions are consistent across modelling approaches and concur with ecological theory. Novel model inspection approaches can reveal details of model predictions, which addresses reasonable concerns about the “black box” nature of many machine learning-based models. We assess model performance and robustness using four datasets encompassing a broad taxonomic range. The number of included species ranges from 20 to 176 and range shifts were observed over time spans ranging from 30 to 100+ years. Each dataset was derived from previous evaluations of traits as range shift predictors and consists of a list of focal species, associated species-level traits, and a range shift metric. We examine (1) whether non-linear methods can improve predictive ability of traits compared to linear methods, (2) whether the novel methods identify important traits consistent with significant results from other studies, and (3) whether the directionality of the modeled effect of traits is consistent across model types.

## 2. MATERIALS AND METHODS

We describe the nonlinear modeling approaches, a framework for interpreting model predictions, and the assembly of range shift and trait data to determine whether traits can play a more powerful role in predicting ecological responses to climate change.

### 2.1 Modeling Approach

We applied three classes of learning algorithms: regularized linear regression, kernel-based regression, and tree-based regression. Regularization is a modification to generalized linear regression that limits model complexity to avoid (Hastie et al., 2009). Several types of regularization exist; we chose to use a “ridge”-regularized linear model, which imposes a penalty on the magnitude of each learned coefficient. The cumulative effect of this regularization procedure is a set of coefficients which both minimize prediction error on the training data and prevent overfitting. These coefficients can be interpreted explicitly as with ordinary least squares regression.

While regularization reduces overfitting when compared to a standard least-squares linear fit, regularized linear models are still not able to capture nonlinearities among or interactions between predictive variables. To remedy this, we employ two additional classes of models: kernel-based regression and tree-based regression. A “kernel” is a function which projects a set of input data, often into a high-dimensional space, to allow for the linear “separability” of the data for the purposes of classification or regression (Hastie et al., 2009). The Kernel Ridge method employed herein uses a radial basis function (or squared exponential) kernel applied to the training data and fits a ridge-regularized linear model to this transformed input. As a result of this transformation the learned coefficients, while regularized, are not immediately interpretable.

We also evaluate a kernel-based technique known as a support vector machine (SVM). This popular learning method can be formulated for regression, is robust to outliers, and can capture nonlinearities and variable interactions through a similar radial basis function kernel as in the Kernel Ridge approach. Like the Kernel Ridge method, the SVM regressor does not have interpretable coefficients. Finally, we train a random forest regression algorithm to evaluate the performance of tree-based methods. All of these models are implemented in the Python programming language using the scikit-learn software package (Pedregosa et al., 2011), though all analyses can be computed in the R language using available machine learning packages. All code for this project is available on GitHub at https://github.com/huckleylab/cc_traits.

For comparison to the original analyses, we train an ordinary least squares regression model, which assumes linear relationships and no variable interaction. While the original analyses were mostly conducted in a single-variable framework (that is, to assess the effect size of *M* different potential predictive variables *M* models were trained, each model containing only 1 variable), we include all variables in single analyses to enable the models to capture variable interactions and to follow a common predictive modeling paradigm. This multivariate approach is a standard one in machine-learning based predictive analytics (Hastie et al., 2009).

### 2.2 Evaluation

We use data subsets for performance evaluation. To assess the predictive performance of our models we employ a *k*-fold cross-validation scheme (Hastie et al. 2009) combined with a squared error loss function. This cross-validation technique has been shown to estimate expected prediction error (Hastie et al. 2009) by randomly partitioning data points into *k* = 10 subsets, each with *N/k* members; *k - 1* subsets are used to fit the model (the “training set”), reserving one subset for testing model performance. Each of the *k* subsets is used exactly once for evaluation. A mean squared error (MSE) loss is then computed for the model prediction of range shift magnitude in the reserved test data, which is the sum of squared differences between predicted and actual range shift magnitude over all data in the testing set. This process is repeated *k* times and a mean statistic is computed across the *k* MSE values that result (known as a “cross-validation mean”). The units of MSE estimates are those of the range shift, so MSE provides a direct assessment of model predictive ability. The same cross-validation mean approach is used to compute average variable importance values to evaluate model drivers, described below. In these experiments we choose *k* = 10, which is generally sufficient for robust performance estimates (Hastie et al. 2009).

### 2.3 Model Interpretation

The core of any basic regression analysis is typically an inspection of the significance and direction of the coefficients of a fitted model. However, the kernel methods employed herein (Kernel Ridge, SVM) do not expose any interpretable coefficients. To address this, we utilize the Shapley additive feature value method, proposed in Lundberg & Lee (2017) and described in Supplementary Material 1. To compare the learning techniques, we use either mean regression coefficients (for OLS and Ridge regression), mean Shapley variable importance values (Kernel Ridge and SVM), or mean Gini variable importance values (Random Forest; Breiman, 2001) to rank all variables such that each feature has an importance ranking for each of the several regression methods. We use the sign of Shapley and coefficient values to compare the directionality of predicted trait drivers for models other than random forests (Gini scores are an unsigned information-theoretic measure). All means are *k*-fold cross-validated means.

### 2.4 Trait and Range Shift Data

We evaluate our approach independently by replicating analysis across four datasets that (1) repeated historical surveys or conducted continuous surveys along latitudinal or elevational gradients to quantify shifts in northern or upper elevation range boundaries over at least three decades of change and (2) included all surveyed species (i.e. rather than including only species that shifted significantly). The first two data sets are those used by Angert et al. (2011) to assess the predictive power of traits. These datasets supplement trait data with elevational range shift data for Swiss alpine plants (Holzinger et al., 2008; *N* = 139) and for Western North American small mammals (Moritz et al., 2008). A third database from Rumpf et al. (2018) consists of elevational range shifts for European montane plants coupled with empirically measured trait data supplemented with data from the TRY Plant Trait Database (Kattge et al., 2011; https://www.try-db.org and Supplementary Information, Section 2.2) and other databases (e.g. Bjorkman et al., 2018). These two plant databases were chosen in an effort to directly replicate data used in previous range shift studies, despite some overlap among them. The fourth database was created by pairing estimates of latitudinal range shifts from coastal North American marine fish surveys (Pinsky et al., 2013) with functional trait data in Fishbase (https://www.fishbase.org, Froese & Pauly, 2010 and Supplementary Information, Section 2.1). Each dataset includes a directional range shift: negative values indicate shifts downward in elevation (m) for the first three datasets and equatorward in latitude (degrees) for the marine dataset. We remove samples which are missing any traits, one-hot encode categorical traits (i.e., generate one boolean column for each category), and normalize/center the numeric traits to have zero mean and unit norm. After this processing the Swiss plants dataset contains *N* = 20 species and *d* = 38 traits (Table S1); the Yosemite mammal dataset contains *N* = 28 species and *d* = 19 traits (Table S2); the European plants dataset contains *N* = 176 species and *d* = 18 traits (Table S3); the marine fish dataset contains *N* = 76 species and *d* = 17 traits (Table S4).

## 3. RESULTS

We find that the machine learning approaches improve predictive performance over an ordinary least squares (OLS) model baseline. For two datasets (Swiss alpine plants and Western NA mammals), the four machine learning approaches perform substantially better than the OLS models but similarly to each other. For the other two datasets (European montane plants and marine fish), the performance advantages of the machine learning approaches are less substantial and the four machine learning approaches differ more in their performance. Support Vector Regression (SVR) and Kernel Ridge emerge as the most performant methods across datasets (Figure 1), reducing mean error in range shift estimates by an average of 62.8% and 61.6% relative to OLS, respectively (medians of 50.6% and 49.6%, respectively). We focus on results from the Swiss alpine plants dataset (Angert et al., 2011; Holzinger et al., 2008) to demonstrate findings. The initial OLS analysis found that individual predictors accounted for relatively little variance in the extent of the plants’ elevational range shifts (R^2^= 0.05-0.18, Angert et al., 2011)

**Figure 1:**
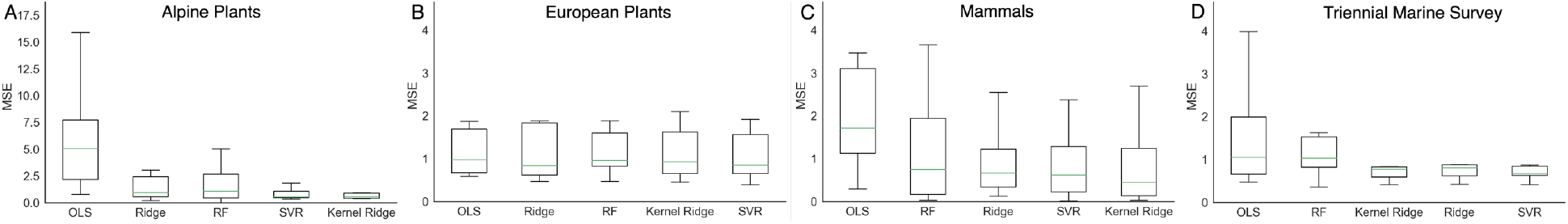
The machine learning approaches reduce the mean squared error (MSE, 10-fold cross-validation) of range shift predictions below the MSE of the standard linear regression approach (OLS: ordinary least squares) across all three of four datasets (A, C, D). Box represents interquartile range (lower to upper), central lines shows data median, and whiskers represent the range of the data. Support Vector Regression (SVR) and Kernel Ridge models exhibit stronger performance than ridge regularized linear (Ridge) or Random Forest (RF) models across the datasets. The MSE units correspond to the range shift metric (A-C: m, D:degrees latitude) and thus directly indicate model performance.

In addition to reducing MSE, the traits found to be important predictors in machine learning models correspond to those identified in previous analyses and ecological theory. The most performant models for these data (SVR and Kernel Ridge, lowest MSE: **Figure 1a)** identify dispersal-related traits (e.g., the timing, height, and duration of seed shed along with seed size and dispersal mode for the Alpine and European Plants) as the most important variables in predicting range shift magnitude **(Figure 2a)**, which is consistent with our expectation and prior work (Angert et al. 2011). In the original analysis of the European Plants data (Rumpf et al., 2018) the indicator of thermal adaptation (cool to warm adaptation, “TemperatureIndicator”) was a primary predictor of range shifts; our models selected the same variable as most important (**Figure 2b**). The previous Western NA mammal analysis (Angert et al. 2011) identified altitudinal limit as a significant predictor and longevity as a relatively strong, but non-significant, predictor. Our models select those two variables as the most important predictors (Figure 2c). Both the initial surveys (Pinsky et al., 2013) and our analysis failed to identify strong trait predictors of marine range shifts and non-linear methods yielded less improvement of MSE than the other datasets. However, habitat traits such as whether species are pelagic are top predictors consistent with compilations of individual studies (Poloczanska et al., 2013).

**Figure 2:**
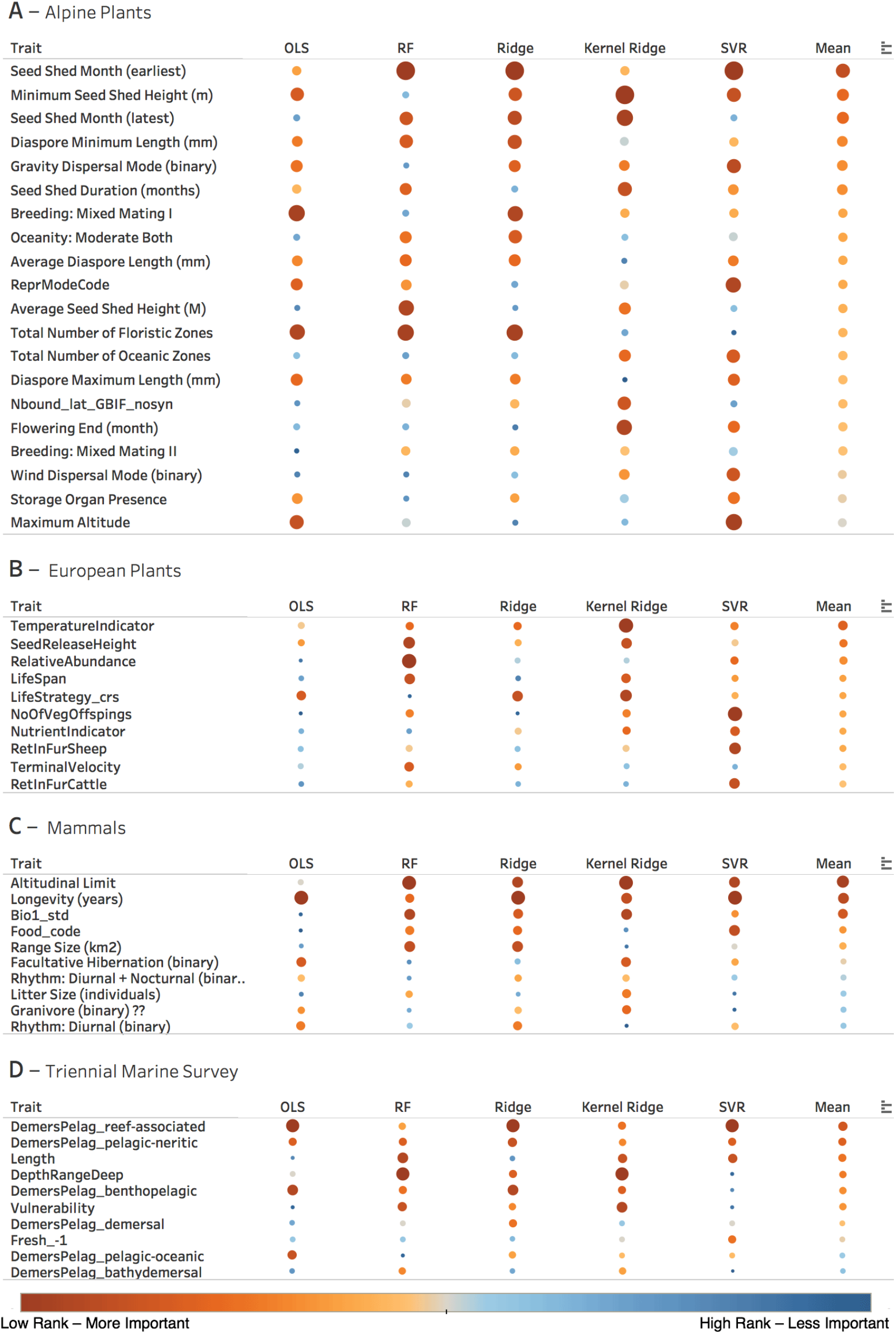
The model approaches select similar traits as important (larger and red = more important) for predicting range shifts across datasets. Traits are listed in order of decreasing mean importance (increasing mean rank, right column) across all methods (except OLS) for each dataset (panels). Model abbreviations are as in Figure 1.

The models employed in this analysis also demonstrate cross-model consistency in identifying trait drivers of range shifts (**Figure 2)**. In other words, separate models tend to agree on trait rank, especially in the top 5 traits, suggesting a common effect despite significantly different modeling methodologies (and evaluation strategies). However, there tends to be more agreement among machine learning models than between OLS and machine learning models. In particular, some traits for which thresholds seem likely are less important predictors in OLS than in machine learning models: higher seed shed heights and longer seed shed durations in alpine plants may not lead to more dispersal once thresholds are reached (Figure 2a).

Finally, models that assign directionality to a modeled effect (all models except Random Forest), tend to agree in the directionality of how traits influence range shifts **(Figure 3)**. For example, all models for European plants (except Kernel Ridge) suggest that thermophilic species that release seeds higher have shifted their altitudinal distribution higher in elevation. All models (except SVR) find that mammals with greater longevity and higher altitudinal limits exhibit smaller altitudinal range shifts. In both these examples, the exceptional model suggests the opposite relationship for both traits. Agreement is sometimes strongest within regression (OLS, Ridge) and machine learning (Kernel Ridge, SVR) type models. For example, the machine learning models suggest alpine plants with higher seed shed shift their distribution further, contrary to the findings of regression models. However, the mixed results suggest limits to the predictive capacity of species’ traits (see Discussion). It is important to note that these measures of variable contribution are computed in distinct ways depending on the modeling methodology, and it remains to be seen whether these methods of variable importance are suitable to determine effect directionality. In addition, using these heterogeneous metrics to compare effect directionality must be done with caution.

**Figure 3:**
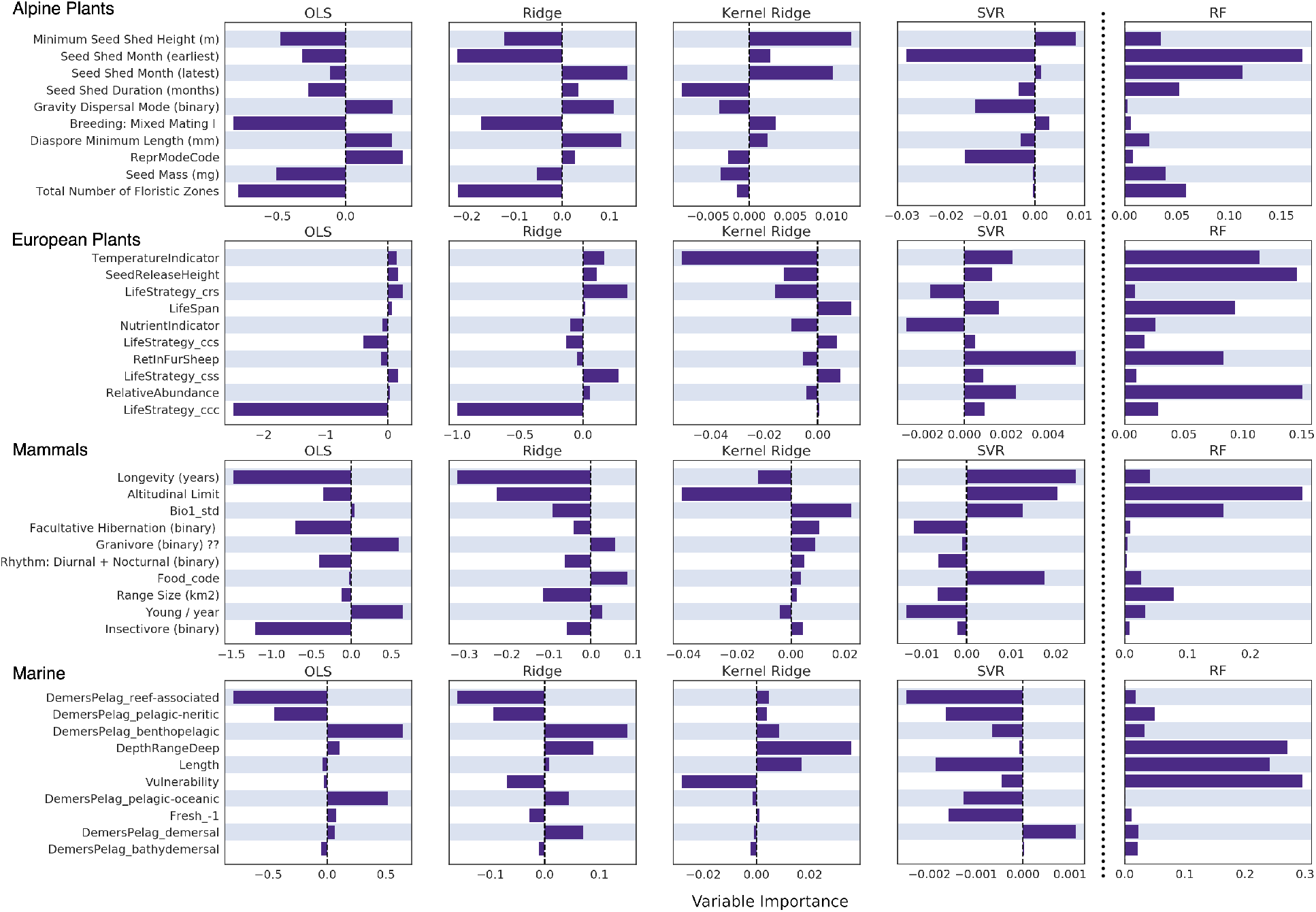
Models tend to agree on the variable importance values and directionality of top-ranking traits. We depict model coefficients (OLS, Ridge), Shapley feature importance values (Kernel Ridge, SVR), or Gini feature importance scores (RF) for top ten traits by rank (Figure [ranks]) for each dataset (rows). Model abbreviations are as in Figure 1 and traits are as in Figure 2.

## 4. DISCUSSION

We find that non-linear modeling methods enhance the ability of traits to accurately predict observed range shifts. Our findings match biological intuition that biological processes, which respond to environmental conditions and are mediated by species’ traits, are rarely linear (Stenseth & Mysterud, 2002). Importantly, we have shown that the novel statistical models maintain biological rigor by identifying similar predictor traits. However, some disagreements in the directionality of the relationship between trait values and range shift magnitude suggest limits to trait-based statistical prediction frameworks.

Overall, our analysis offers a mixed outlook for using species’ traits in applied predictions, such as analyses of climate change vulnerability. The substantially better predictive performance of non-linear models relative to linear models suggests vulnerability analyses frameworks based on species’ traits (e.g., Foden et al., 2013; Pacifici et al., 2017) should be adapted to account for non-linearities. However, disagreements in the directionality of trait predictors and the variability of performance improvement suggest that even non-linear methods for relating traits to climate change responses may have limited predictive accuracy. More mechanistic approaches that describe the processes by which traits mediate fitness and demographic responses to the environment may be required for predictions that require high levels of accuracy (Buckley & Kingsolver, 2012; Urban et al., 2016). At a minimum, these mechanistic approaches will be useful for refining non-linear methodologies for using traits to predict climate change responses.

Models that allow for non-linearities should be employed to further reevaluate expectations for how traits govern range shifts. A meta-analysis across range shift studies found at best moderate support for dispersal ability (body size: 22%, migratory strategy: 10%, movement ability: 50% of studies uphold predicted relationship), reproductive potential (fecundity: 36%, longevity: 60%) and ecological generalization (diet breadth: 27%, habitat breadth: 43%) as predictors of range shift magnitude (MacLean & Beissinger, 2017). The large gap between expectations and observations highlights the need for novel predictive methods. Translating spatial range shifts into metrics of environmental niche tracking (e.g., velocity of climate change, Loarie et al., 2009) may also enhance predictive capacity.

Reevaluation with non-linear methods will provide insight into selecting appropriate predictor traits. The availability of trait data has increased substantially since some of our datasets were compiled (e.g., Angert et. al 2011), so refining traits may improve predictive capacity. Still, needs for additional trait data addressing issues such as physiology and evolutionary potential are substantial and will likely require concerted data collection efforts (Urban et al., 2016). Since species’ traits are likely to be phylogenetically conserved, phylogenetic signal in range shifts can be used to assess the potential to use traits to predict range shifts. High phylogenetic signal but weak predictive performance of traits would suggest that improving the traits used as predictors can enhance predictive capacity. The initial analyses (e.g., Angert et. al 2011) that accounted for phylogeny found limited phylogenetic signal in range shifts. We did not account for phylogeny because it is not straightforward to do so in the machine learning models. A recent synthesis of range shift studies (Diamond, 2018) found variable but generally weaker phylogenetic signal in range shifts than in physiological, morphological, and life-history traits. The finding indicates limits to the predictive capacity of traits.

Our methodology—flexible statistical models paired with robust evaluation methods and model interrogation approaches—has been reliably employed across many predictive contexts but has yet to be fully embraced by ecologists. The emerging wealth of publicly-available ecological and environmental data, combined with the pressing need for reliable ecological forecasts that are useful in decision-making frameworks, makes this flexible and data-intensive approach a natural fit. Despite promising results, our approach presents several challenges to adoption. Of particular relevance to the ecology community is the lack of traditional statistical techniques to evaluate these methods. Following the machine learning community, we employ *k*-fold cross-validation to lend statistical robustness to the pertinent evaluative criteria for our models (here, mean squared error). In addition, the use of recent advances in model inspection methods (Shapley values from Lundberg & Lee, 2017) represents a necessary departure from the manual inspection and testing of linear model coefficients. As the field of model interpretation grows, ecologists can leverage these developments to verify the ecological processes underpinning the predictions of these unconventional modeling approaches. Understanding and acknowledging these shifts in method evaluation and inspection approaches are critical steps to leveraging these more performant statistical modeling approaches in the ecology community.

The general applicability of this approach should be confirmed by studies including additional taxa, a greater number of species, and using novel model interrogation techniques. In addition, the approach has the potential to improve predictive accuracy in other ecological domains relevant to policy and decisions making (species distribution modeling, forecasting of ecological carbon flux, etc.). However, as is demonstrated by the range of predictive performance improvement across datasets (e.g. between the Alpine Plants and the European Plants datasets, Figure 1), care must be taken to evaluate any novel modeling result conservatively and through a lens of data quality and available latitude for improvement.

## ACKNOWLEDGEMENTS

We thank contributors to TRY, Fishbase, and the other datasets and input and assistance from Amy Angert, Malin Pinsky, Ray Huey, and particularly Sabine Rumpf. This work was supported by the National Science Foundation [IGERT DGE-1258485 fellowship to A.F.C., a Graduate Research Fellowship to A.F.C, and DBI-1349865 to L.B.B.].

## References

Adrian, R., Wilhelm, S., & Gerten, D. (2006). Life-history traits of lake plankton species may govern their phenological response to climate warming. Global Change Biology, 12(4), 652–661.

Altermatt, F. (2010). Tell me what you eat and I’ll tell you when you fly: diet can predict phenological changes in response to climate change. Ecology Letters, 13(12), 1475–1484.

Angert, A. L., Crozier, L. G., Rissler, L. J., Gilman, S. E., Tewksbury, J. J., & Chunco, A. J. (2011). Do species’ traits predict recent shifts at expanding range edges? Ecology Letters, 14(7), 677–689.

Bjorkman, A. D., Myers-Smith, I. H., Elmendorf, S. C., Normand, S., Thomas, H. J. D., Alatalo, J. M.,…Zamin, T. (2018). Tundra Trait Team: A database of plant traits spanning the tundra biome. Global Ecology and Biogeography, 27(12), 1402–1411.

Breiman, L. (2001). Random Forests. Machine Learning, 45(1), 5–32.

Buckley, L. B., & Kingsolver, J. G. (2012). Functional and Phylogenetic Approaches to Forecasting Species’ Responses to Climate Change. Annual Review of Ecology, Evolution, and Systematics, 43(1), 205–226.

Diamond, S. E., Frame, A. M., Martin, R. A., & Buckley, L. B. (2011). Species’ traits predict phenological responses to climate change in butterflies. Ecology, 92(5), 1005–1012.

Estrada, A., Morales-Castilla, I., Caplat, P., & Early, R. (2016). Usefulness of Species Traits in Predicting Range Shifts. Trends in Ecology & Evolution, 31(3), 190–203.

Fitt, R. N. L., Palmer, S., Hand, C., Travis, J. M. J., & Lancaster, L. T. (2018). Towards an interactive, process-based approach to understanding range shifts: developmental and environmental dependencies matter. Ecography, 0(0).

Foden, W. B., Butchart, S. H. M., Stuart, S. N., Vié, J.-C., Akçakaya, H. R., Angulo, A.,…Mace, G. M. (2013). Identifying the World’s Most Climate Change Vulnerable Species: A Systematic Trait-Based Assessment of all Birds, Amphibians and Corals. PLOS ONE, 8(6), 1–13.

Froese, R., & Pauly, D. (2010). FishBase.

Hastie, T., Tibshirani, R., & Friedman, J. (2009). The Elements of Statistical Learning. In The Mathematical Intelligencer.

Holzinger, B., Hülber, K., Camenisch, M., & Grabherr, G. (2008). Changes in plant species richness over the last century in the eastern Swiss Alps: elevational gradient, bedrock effects and migration rates. Plant Ecology, 195(2), 179–196.

Kattge, J., Díaz, S., Lavorel, S., Prentice, I. C., Leadley, P., Bönisch, G.,…Wirth, C. (2011). TRY - a global database of plant traits. Global Change Biology, 17(9), 2905–2935.

Lundberg, S. M., & Lee, S.-I. (2017). A Unified Approach to Interpreting Model Predictions. In Guyon, U. V Luxburg, S. Bengio, H. Wallach, R. Fergus, S. Vishwanathan, & R. Garnett (Eds.), Advances in Neural Information Processing Systems 30 (pp. 4765–4774).

MacLean, S. A., & Beissinger, S. R. (2017). Species’ traits as predictors of range shifts under contemporary climate change: A review and meta-analysis. Global Change Biology, 23(10), 4094–4105.

Maguire, K. C., Nieto-Lugilde, D., Fitzpatrick, M. C., Williams, J. W., & Blois, J. L. (2015). Modeling Species and Community Responses to Past, Present, and Future Episodes of Climatic and Ecological Change. Annual Review of Ecology, Evolution, and Systematics, 46(1), 343–368.

Moritz, C., Patton, J. L., Conroy, C. J., Parra, J. L., White, G. C., & Beissinger, S. R. (2008). Impact of a Century of Climate Change on Small-Mammal Communities in Yosemite National Park, USA. Science, 322(5899), 261–264.

Pacifici, M., Visconti, P., Butchart, S. H. M., Watson, J. E. M., Cassola, F. M., & Rondinini, C. (2017). Species’ traits influenced their response to recent climate change. Nature Climate Change, 7(3), 205–208.

Parmesan, C. (2006). Ecological and Evolutionary Responses to Recent Climate Change. Annual Review of Ecology, Evolution, and Systematics, 37(1), 637–669.

Pedregosa, F., Varoquaux, G., Gramfort, A., Michel, V., Thirion, B., Grisel, O.,…Duchesnay, E. (2011). Scikit-learn: Machine Learning in Python. Journal of Machine Learning Research, 12, 2825–2830.

Pinsky, M. L., Worm, B., Fogarty, M. J., Sarmiento, J. L., & Levin, S. A. (2013). Marine Taxa Track Local Climate Velocities. Science, 341(6151), 1239–1242.

Poloczanska, E. S., Brown, C. J., Sydeman, W. J., Kiessling, W., Schoeman, D. S., Moore, P. J.,…Richardson, A. J. (2013). Global imprint of climate change on marine life. Nature Climate Change, 3, 919.

Rapacciuolo, G., Maher, S. P., Schneider, A. C., Hammond, T. T., Jabis, M. D., Walsh, R. E.,…Beissinger, S. R. (2014). Beyond a warming fingerprint: individualistic biogeographic responses to heterogeneous climate change in California. Global Change Biology, 20(9), 2841–2855.

Rumpf, S. B., Hülber, K., Klonner, G., Moser, D., Schütz, M., Wessely, J.,…Dullinger, S. (2018). Range dynamics of mountain plants decrease with elevation. Proceedings of the National Academy of Sciences, 115(8), 1848–1853.

Schloss, C. A., Nuñez, T. A., & Lawler, J. J. (2012). Dispersal will limit ability of mammals to track climate change in the Western Hemisphere. Proceedings of the National Academy of Sciences, 109(22), 8606–8611.

Stenseth, N. C., & Mysterud, A. (2002). Climate, changing phenology, and other life history traits: Nonlinearity and match-mismatch to the environment. Proceedings of the National Academy of Sciences, 99(21), 13379–13381.

Urban, M. C., Bocedi, G., Hendry, A. P., Mihoub, J.-B., Pe’er, G., Singer, A.,…Travis, J. M. J. (2016). Improving the forecast for biodiversity under climate change. Science, 353(6304).

Wheatley, C. J., Beale, C. M., Bradbury, R. B., Pearce-Higgins, J. W., Critchlow, R., & Thomas, C. D. (2017). Climate change vulnerability for species-Assessing the assessments. Global Change Biology, 23(9), 3704–3715.

